# Variant effect predictor correlation with functional assays is reflective of clinical classification performance

**DOI:** 10.1101/2024.05.12.593741

**Authors:** Benjamin J. Livesey, Joseph A. Marsh

## Abstract

**Background:** Understanding the relationship between protein sequence and function is crucial for accurate genetic variant classification. Variant effect predictors (VEPs) play a vital role in deciphering this complex relationship, yet evaluating their performance remains challenging for several reasons including data circularity, where the same or related data is used for training and assessment. High-throughput experimental strategies like deep mutational scanning (DMS) offer a promising solution.

**Results:** In this study, we extend upon our previous benchmarking approach, assessing the performance of 97 different VEPs using DMS experiments from 36 different human proteins. In addition, a new pairwise, VEP-centric ranking method reduces the impact of missing predictions on the overall ranking. We observe a remarkably high correspondence between VEP performance in DMS-based benchmarks and clinical variant classification, especially for predictors that have not been directly trained on human clinical variants.

**Conclusions:** Our results suggest that comparing VEP performance against diverse functional assays represents a reliable strategy for assessing their relative performance in clinical variant classification. However, major challenges in clinical interpretation of VEP scores persist, highlighting the need for further research to fully leverage computational predictors for genetic diagnosis. We also address practical considerations for end users in terms of choice of methodology.

## Background

Deciphering the nature of the sequence-function relationship in proteins remains one of the greatest challenges in modern biology. It has profound implications for variant classification in a medical context, understanding of disease mechanisms and protein design. Computational tools for predicting variant effects, known as variant effect predictors (VEPs), can provide valuable insight into the complex relationship between protein sequence and human phenotypes. However, the profusion of new predictors has also highlighted the need for reliable, unbiased strategies for evaluating VEP performance.

One of the main obstacles to identifying a fair method for comparing VEPs is the prevalence of data circularity in many performance evaluations [1]. This often results in an inflated assessment of VEP performance and can be introduced into a benchmark in two ways. Variant-level circularity, often referred to as ‘Type 1’, occurs when specific variants (or homologous variants) used to train or tune a VEP are subsequently used to assess its performance. Gene-level circularity, often referred to as ‘Type 2’, occurs in cross-gene analyses when the testing set contains different variants from the same (or homologous) genes used for training. It arises because predictors learn associations between different genes and pathogenicity. For example, if a VEP learns to strongly associate variants from a specific gene as mostly being pathogenic or benign, this can lead to excellent apparent performance if the tested variants from this gene mostly fall into the same class.

Both variant- and gene-level circularities can be difficult to address, and doing so often greatly reduces the pool of available data to compare the performance of predictors based upon supervised learning approaches, or reduces the number of VEPs that can be compared in order to expand the pool of variants. Even amongst predictors that have not been directly trained on clinical labels, some have been exposed to human population variants through direct inclusion of allele frequency as a feature, or through indirect tuning. Given that allele frequency is routinely used as strong evidence to classify variants as benign [2], VEPs that utilise information on human population variants may have effectively ‘seen’ a large proportion of the benign data used to benchmark them. These limitations keep most independent benchmarks at a small scale, and often limited to comparing less than 10 different VEPs [3–5].

One way to address this problem came from the development of high-throughput experimental strategies, known as multiplexed assays of variant effect (MAVEs) [6]. The technology behind MAVEs has been improving at a rapid rate, helped in part by the Atlas of Variant Effects Alliance, which aims to promote research and collaboration, and eventually produce variant effect maps across all human protein-coding genes and regulatory elements [7]. Datasets derived from deep mutational scanning (DMS) experiments, a class of MAVEs focusing on functional assays for measuring the effects of protein mutations [8], show tremendous potential for use as a baseline for comparing the outputs of VEPs. DMS datasets provide several advantages for benchmarking over more traditional sets composed of variants observed in a clinical context. They do not rely upon any previously assigned clinical labels (*e.g.* ‘pathogenic’ and ‘benign’) that are commonly used to train VEPs, thus greatly reducing the potential for variant-level (type 1) circularity in any assessment of VEP performance. By comparing the Spearman’s correlations between variant effect scores from VEPs and DMS experiments on a per-protein basis, gene-level (type 2) circularity is also avoided. However, the downside of using DMS data for this purpose is that the specific functional assay employed in each DMS study may not be relevant to any disease mechanisms for that particular protein.

Previous work has used DMS datasets to investigate VEP performance [9]. For example, the Critical assessment of Genome Interpretation (CAGI) community experiment [10] has used unreleased DMS datasets as challenges, comparing the agreement of certain missense VEPs with functional assays from individual proteins, including UBE2I [11], PTEN [12] and HMBS [13]. ProteinGym contains a collection of DMS datasets aimed at comparing the ability of different machine learning models to predict mutational effects [14]. We have also evaluated the performance of VEPs [15,16] and protein stability predictors [17] using data from DMS assays. With the rapid recent growth in the field, numerous novel VEPs and DMS datasets have been subsequently released. Here we build upon this work by including 43 more VEPs and 13 more human DMS datasets, and also by improving our benchmarking methodology. Our work demonstrates an extremely high correspondence between VEP performance when benchmarked against DMS datasets and when tested for clinical variant classification when we consider those predictors that have not been directly trained on human clinical or population variants. In contrast, for VEPs trained or tuned on human variants, it is exceedingly difficult to perform a fair comparison using traditional clinical benchmarks. Therefore, we suggest that our strategy of benchmarking VEPs using numerous diverse DMS datasets represents a reliable way of assessing their relative performance at scoring the clinical impacts of variants within individual proteins. Importantly, however, we acknowledge that considerable challenges remain in fully interpreting VEP outputs for clinical applications.

## Results

### New DMS datasets and VEPs

The increasing popularity of MAVEs as an experimental strategy for high-throughput characterisation of variant effects has enabled us to add 13 new DMS datasets assessing the impacts of single amino acid substitutions to our benchmark compared to our previous publications (Table 1). Many of these are present in MaveDB, a valuable community resource for the sharing of MAVE datasets [18]. As each DMS dataset can have multiple sets of functional scores, potentially representing altered conditions, replicates, filters or even entirely different fitness assays, we chose a single DMS score set per protein to represent the overall fitness of that protein. We selected the dataset that had the highest median Spearman’s correlation with all VEPs in order to prevent outliers from overtly influencing the selection process. For proteins in which multiple DMS studies were performed by different groups (*TP53*, *GDI1*, *PTEN*), we likewise only selected a single score set for each protein using the above method. We also require a minimum of 1000 single amino acid substitutions to be scored (following the removal of variants in ClinVar [19] and gnomAD [20]) in order to prevent low-coverage DMS assays from influencing the outcome. Furthermore, we excluded DMS assays that assessed antibody binding (*CXCR4, CD19, CCR5*) and affinity for other binding targets unrelated to the normal biological function of the protein (*ACE2* to SARS-CoV-2 receptor binding domain) as well as those without any associated methodological details (*NCS1, TECR*). The full summary of all DMS datasets, across 36 proteins and covering 207,460 different single amino acid variants, is available in Table S1, additional file 1.

**Table 1.**
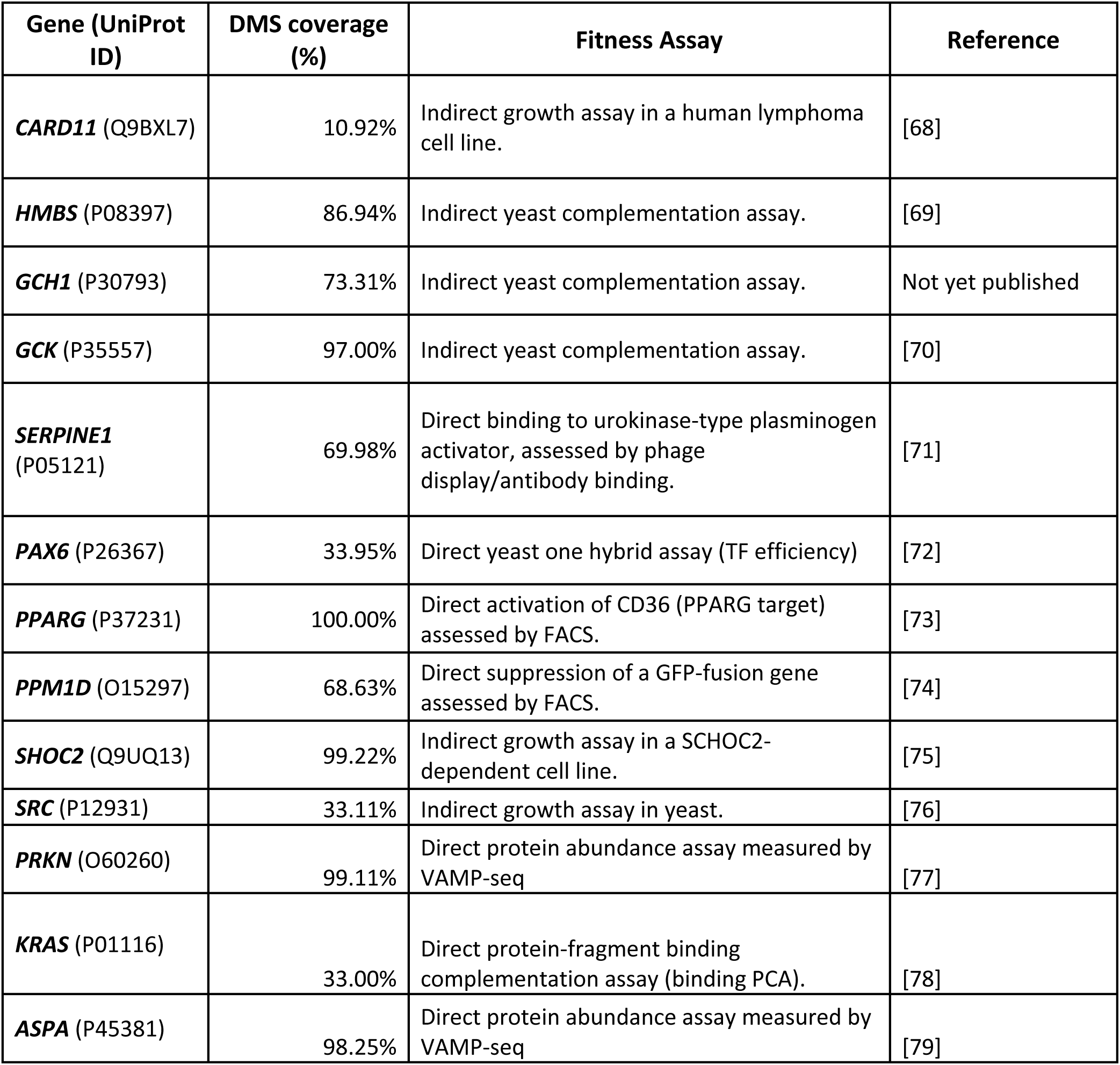
A summary of the 13 new DMS datasets included in this benchmark.

| Gene (UniProt ID) | DMS coverage (%) | Fitness Assay | Reference |
| --- | --- | --- | --- |
| **CARD11** (Q9BXL7) | 10.92% | Indirect growth assay in a human lymphoma cell line. | [68] |
| **HMBS** (P08397) | 86.94% | Indirect yeast complementation assay. | [69] |
| **GCH1** (P30793) | 73.31% | Indirect yeast complementation assay. | Not yet published |
| **GCK** (P35557) | 97.00% | Indirect yeast complementation assay. | [70] |
| **SERPINE1** (P05121) | 69.98% | Direct binding to urokinase-type plasminogen activator, assessed by phage display/antibody binding. | [71] |
| **PAX6** (P26367) | 33.95% | Direct yeast one hybrid assay (TF efficiency) | [72] |
| **PPARG** (P37231) | 100.00% | Direct activation of CD36 (PPARG target) assessed by FACS. | [73] |
| **PPM1D** (O15297) | 68.63% | Direct suppression of a GFP-fusion gene assessed by FACS. | [74] |
| **SHOC2** (Q9UQ13) | 99.22% | Indirect growth assay in a SCHOC2-dependent cell line. | [75] |
| **SRC** (P12931) | 33.11% | Indirect growth assay in yeast. | [76] |
| **PRKN** (O60260) | 99.11% | Direct protein abundance assay measured by VAMP-seq | [77] |
| **KRAS** (P01116) | 33.00% | Direct protein-fragment binding complementation assay (binding PCA). | [78] |
| **ASPA** (P45381) | 98.25% | Direct protein abundance assay measured by VAMP-seq | [79] |

We streamlined the assignment of categories to DMS datasets by defining only two different types. Direct assays are those that directly measure the ability of the target protein to carry out one or more functions. Examples of direct functional assays include one-hybrid and two-hybrid assays, other assays that measure the interaction strength with native partners and VAMP-seq [21]. Indirect assays are most commonly growth rate experiments, where the attribute being measured is not directly controlled by the target protein. Indirect DMS assays may be more representative of the biological reality of a variant’s effect on cellular fitness, as the cell may be able to buffer a small or moderate loss of function. Direct DMS assays are more reflective of a protein’s function in isolation and may be more useful for exploring protein mechanisms or for protein design.

The field of variant effect prediction has also been progressing rapidly, with many novel methods being published every year. In total, we added 43 new missense VEPs that were not used in our previous benchmarks (Table 2). These were identified by browsing new publications, from the Variant Impact Predictor database (VIPdb) [22], and from the ProteinGym resource, which benchmarks numerous VEPs and general protein language models against human and non-human DMS datasets [14]. The large majority of VEPs from our previous analysis [16] were retained, although a small number were removed because the predictors were no longer available to run (NetDiseaseSNP, PonPS and PAPI), and thus we could not add predictions for the new DMS datasets. This emphasises the importance of making source code and pre-calculated variant effect scores freely available, to ensure that tools can continue to be used in the future [23]. In total, we included 97 different VEPs in this study, considering only those with predictions available for at least 50% of the DMS datasets in our benchmark (Table S2, additional file 1).

**Table 2.**
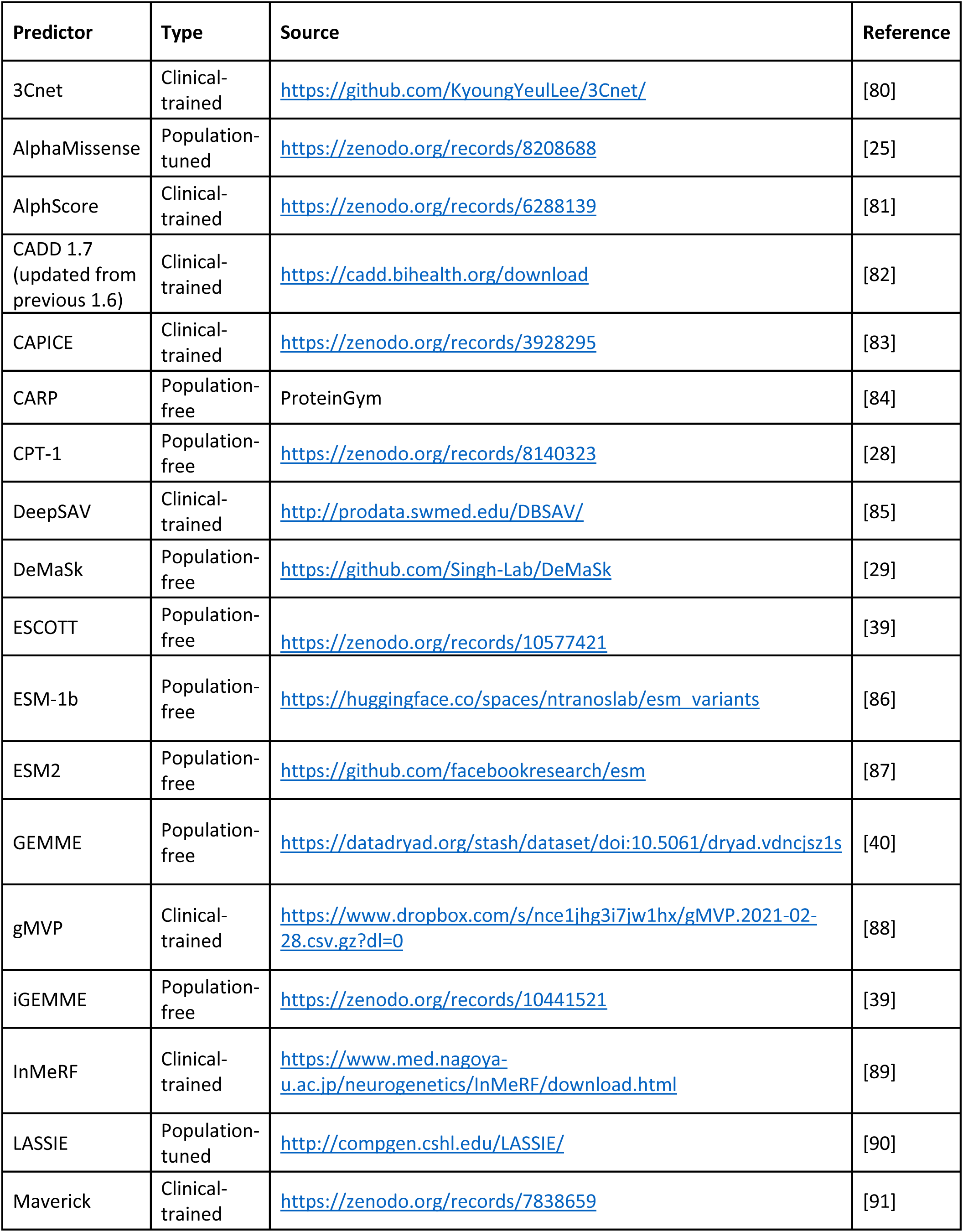
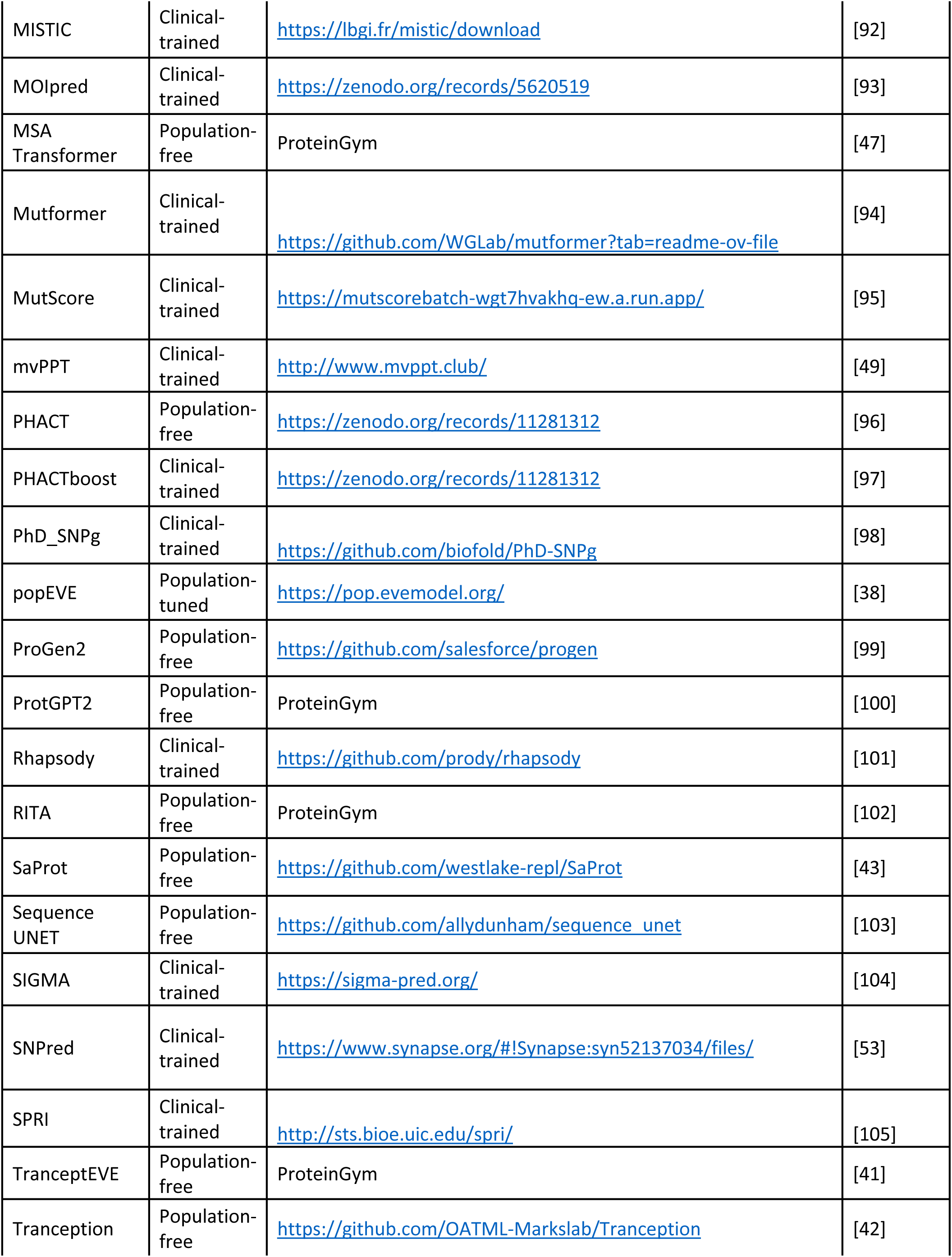
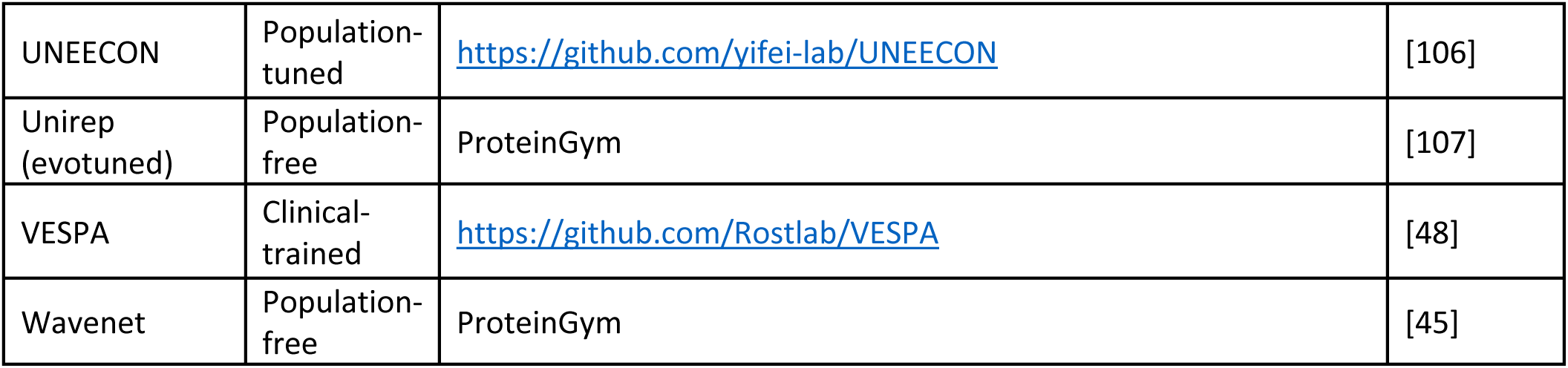
A summary of the 43 new VEPs including in this analysis and links to the data or source code used to produce predictions.

During our research we also identified some VEPs that were difficult to access due to a requirement for paid subscriptions or restrictive licensing agreements. We have not included these in our benchmark, as we strongly believe that, if VEPs are going to be used as stronger evidence when making clinical diagnoses, their methodologies and predictions need to be made freely available to enable fair, independent, replicable assessment by the community [23].

Our previous benchmark classified VEPs into two groups: “supervised” and “unsupervised”. However, we found these categories imperfect for our primary concern, which is the risk of data circularity. For example, Envision [24] is trained by supervised machine learning using DMS data, but has not seen any labelled human variants. Likewise, AlphaMissense is primarily an unsupervised method, but undergoes training with human allele frequencies as “weak labels” [25], which could potentially provide an advantage for classification of benign variants. To address this issue, we have now classified VEPs into three groups that better reflect the risk of data circularity when being assessed using human variants:

- *Clinical-trained* predictors have been trained using human variants with clinical labels (*i.e.* pathogenic or benign), typically derived from databases like ClinVar or HGMD [26]. Most supervised machine learning methods fall into this category. This group also includes methods that, while not directly trained on pathogenic or benign variants, include another predictor that does as a feature (*e.g.* we classify CADD as clinical-trained because it uses PolyPhen-2, which was directly trained on clinical labels). This category is most at-risk of data circularity or bias when assessed using clinical or population data [27], and includes 53 of the VEPs tested in this study.
- *Population-tuned* predictors are not directly trained on clinical variants, but they have been exposed to human population data via tuning, optimisation or scaling processes. This group has a much smaller risk of data circularity, but it still exists, especially if the methods use allele frequency. This is a small group, comprising only six of the VEPs tested here, including AlphaMissense.
- *Population-free* predictors are not trained using any human population data, and thus should be at no risk of data circularity if assessed on clinical or population variants. This category mostly overlaps with what we have previously referred to as “unsupervised”, and includes protein language models and models based on sequence alignments. This group includes 38 of the VEPs used in this study.

Related to this, a recent strategy that has the potential to confound our benchmarking methodology is the increasing availability of predictors that are directly trained using DMS data. There are currently five such VEPs included in our benchmark: Envision, CPT-1 [28], DeMaSk [29], VARITY_R and VARITY_ER [30]. Using DMS datasets to benchmark these VEPs carries similar caveats to benchmarking clinical-trained VEPs using population databases. Fortunately, these methods have all been trained using only a small number of DMS datasets. Therefore, by excluding the results of these VEPs for the proteins used to train them (Table S2, additional file 1), we have been able to include these predictors in our benchmark.

### Comparison of VEP performance using DMS data

Although there is great diversity in DMS assays, what they measure may not always be reflective of clinical outcomes, the premise of our analysis is that VEPs that show the most consistency across a large set of DMS experiments are likely to be the most useful for predicting variant effects. We use absolute Spearman’s rank correlation to assess the correspondence between VEPs and DMS, as only a monotonic relationship between the two variables is required. Thus, no transformation needs to be applied to the VEP output or DMS datasets, which can vary greatly in scale and directionality. The ability of DMS to score large numbers of variants also allowed us to exclude all variants present in ClinVar and gnomAD from calculations of Spearman’s correlation in order to help offset any advantage gained by certain VEPs against the data we use to assess their performance. While these data do not exhaustively cover all VEP training data sources, they are by far the most common databases trained against and certainly have high degrees of overlap with the remaining training data.

Figure 1 shows the Spearman’s correlation between DMS results and VEP predictions for each of the 36 DMS datasets. The strongest correlations approached ρ=0.8 for some DMS datasets (*GCH1*, *PPARG*, *GDI1*), while several others demonstrated relatively poor correlations even for the best-performing VEPs around ρ = 0.3 (*LDLRAP1*, *TPK1*). The average correlation of the top-performing predictors for each protein was 0.58. The population-free predictors achieve top correlations with 20 of the 36 DMS datasets, while population-tuned methods come top on a further 9 datasets (a trend driven almost exclusively by AlphaMissense). The VEP category with the least number of top-correlations with DMS is the clinical-trained category, coming top on only 7 datasets. While VEP performance is protein dependent to some extent, specific VEPs clearly have a higher level of consistency against DMS data, specifically AlphaMissense and CPT-1. The level of heterogeneity in VEP performance is illustrated in Figure S1, where the distribution of Spearman’s correlations for each VEP across all DMS datasets where it has predictions is shown. There was also little difference between the direct and indirect DMS categories: the average correlation of the top-performing VEPs was ρ = 0.61 for the direct assays compared to ρ = 0.56 for the indirect assays.

**Figure 1.**
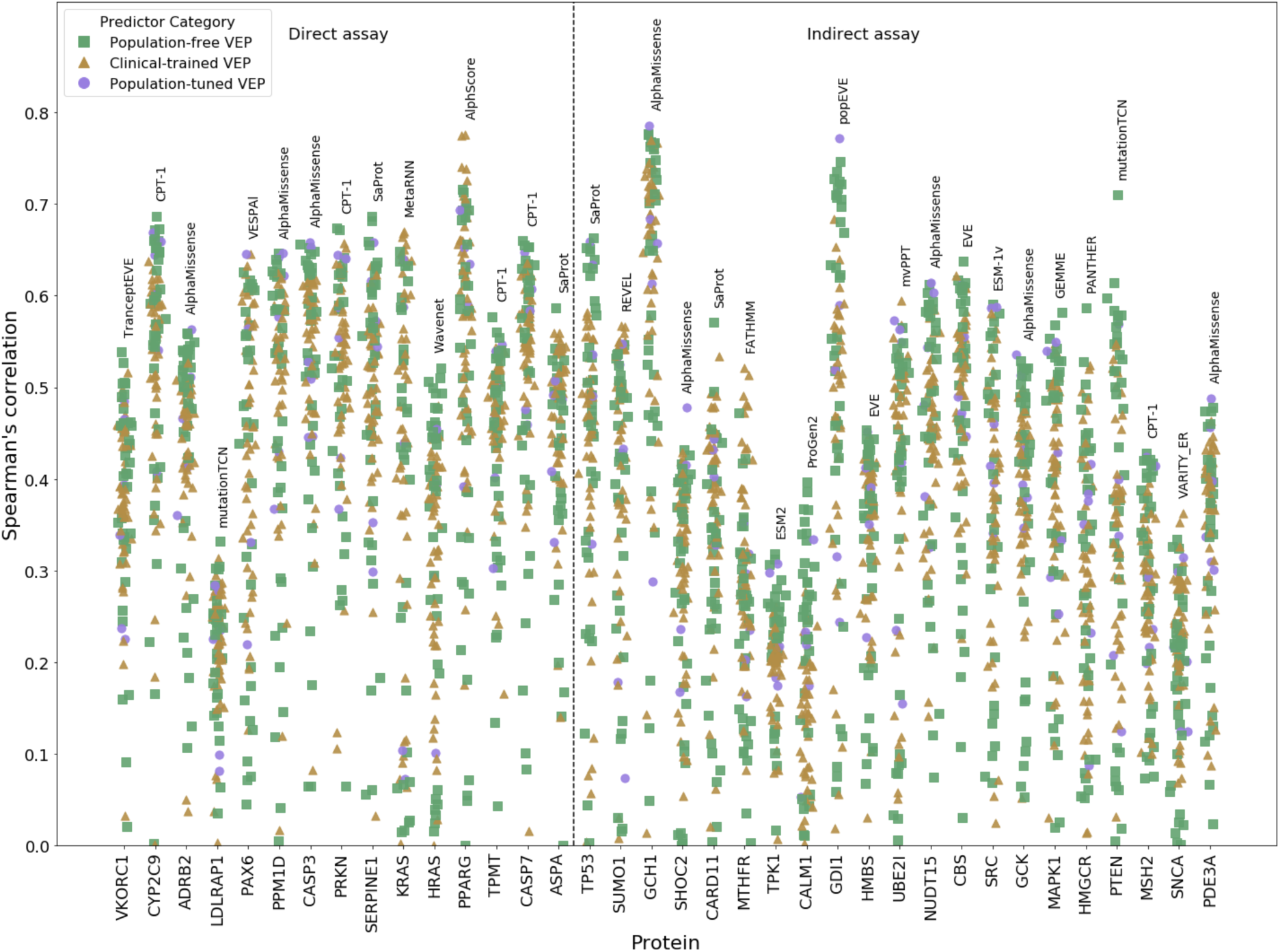
Correlation between variant effect scores from VEPs and DMS experiments. The Spearman’s correlation between all VEPs and every selected DMS dataset. VEPs are split into ‘population-based’ and ‘clinical-trained’ and ‘population-tuned’ methods based on the usage of human clinical and population variants during training. DMS datasets are classified as ‘direct’ if they directly measure the ability of the target protein to carry out one or more functions, with all others being classified as ‘indirect’. The VEP with the highest correlation is noted for every DMS dataset.

Comparing VEPs using correlation to DMS datasets is now commonly used both in papers presenting new prediction methods and in independent benchmarks of predictor performance [14,31]. There are, however two major limitations associated with using raw DMS data in the context of large-scale benchmarking VEPs. First, not all VEPs have predictions available across all proteins. In our case, we were sometimes limited by the specific proteins for which the authors have provided pre-calculated results, or the proteins for which predictions are available for in ProteinGym. In other cases, as mentioned above, we have excluded specific proteins from the assessment of predictors that were trained using DMS data, to avoid potential data circularity.

Second, not all VEPs output scores for all possible variants in the proteins for which they are run. Some only provide predictions for those missense changes possible via single nucleotide changes, while others do not provide predictions for specific protein regions, for example, where the sequence alignment depth is low [32], or when the method can only be applied to sequences shorter than a specific size [31]. This could lead to inflated results for predictors that exclude a generally poorly predicted region of a protein. To illustrate the potential impact of this, we can observe in Figure 1, a few examples where a single outlier VEP demonstrated far higher correlations with the DMS data than other methods, such as PANTHER [33] for *HMGCR* and mutationTCN [34] for *PTEN*. Closer inspection reveals that, in both of these cases, the phenomenon arises because these VEPs provide scores for a much smaller set of variants for these specific proteins than other VEPs.

The varying coverage levels of VEPs for the DMS scores in our dataset are shown in Figure S2. In general, VEPs can be approximately divided into three groups in terms of the proportion of total variants they can provide predictions for. Predictors that provide full or near-full coverage are largely population-free methods, as these are more likely to have been applied to all possible single amino acid substitutions. Another group comprising roughly half of the VEPs includes those with coverage for ∼30% of possible amino acid variants. These are mostly clinical-trained VEPs and the reason for their lower coverage is that they have been applied only to actual missense variants, *i.e.* single amino acid substitutions possible via single nucleotide variants (SNVs). It should also be noted that these predictors are not necessarily restricted to SNVs (such as PolyPhen-2 [35]), but the predictions available for mass download were mapped to SNVs. Finally, several VEPs show varying degrees of intermediate coverage, which can be due to a number of reasons, including data availability, mapping issues and low MSA coverage.

One solution to the problem of varying coverage would be to only use DMS measurements for variants with scores available across all predictors, although this would require us to include either far fewer VEPs (particularly excluding those that fail to provide predictions for one or more proteins) or far fewer DMS sets to retain enough data. Moreover, it is not clear that Spearman’s correlations between VEP and DMS scores should be comparable between proteins, or that they represent a good measure of the relative performance of VEPs across different proteins. For example, a protein with two functionally distinct regions, where one is highly constrained (*e.g*. a globular domain) and the other is not (*e.g.* a disordered region) might show a high correlation between VEPs and DMS, driven by this difference. In contrast, a small, highly conserved protein where mutations at most positions will have damaging effects might show a much lower Spearman’s correlation, even though VEPs are not necessarily performing worse.

Therefore, to ensure that the relative ranking of VEP performance remains as fair as possible, for each DMS dataset we perform a series of pairwise comparisons in which the correlation between every possible pair of VEPs with the DMS data is calculated using only predictions for variants shared between the two VEPs and the DMS set. The percentage of the time that a VEP “won” each of its pairwise comparisons against every other VEP is then calculated across all proteins. This strategy is illustrated in Figure S3, which shows a heatmap coloured by the win rate of each VEP compared to all others. To obtain our overall ranking, we simply average the win rate of each VEP against all other VEPs. This method of ranking is more VEP-centric than DMS-centric as in our previous benchmarks, meaning it should act as a more useful basis for relative ranking, particularly when accounting for cases where certain VEPs do not have predictions for all proteins.

Figure 2 shows the average win rate of the top 30 predictors. The full results, including the win percentage of each VEP against every other, are available in Table S3, additional file 1. The order of predictors in Figure S3 is also sorted according to this same ranking, allowing for visualisation of performance across all predictors. The ranking of predictors is also relatively robust to data permutations, the error bars of Figure 2 represent the standard deviation in the rank score over 1000 bootstraps of the analysis with 36 randomly selected protein datasets (with replacement).

**Figure 2.**
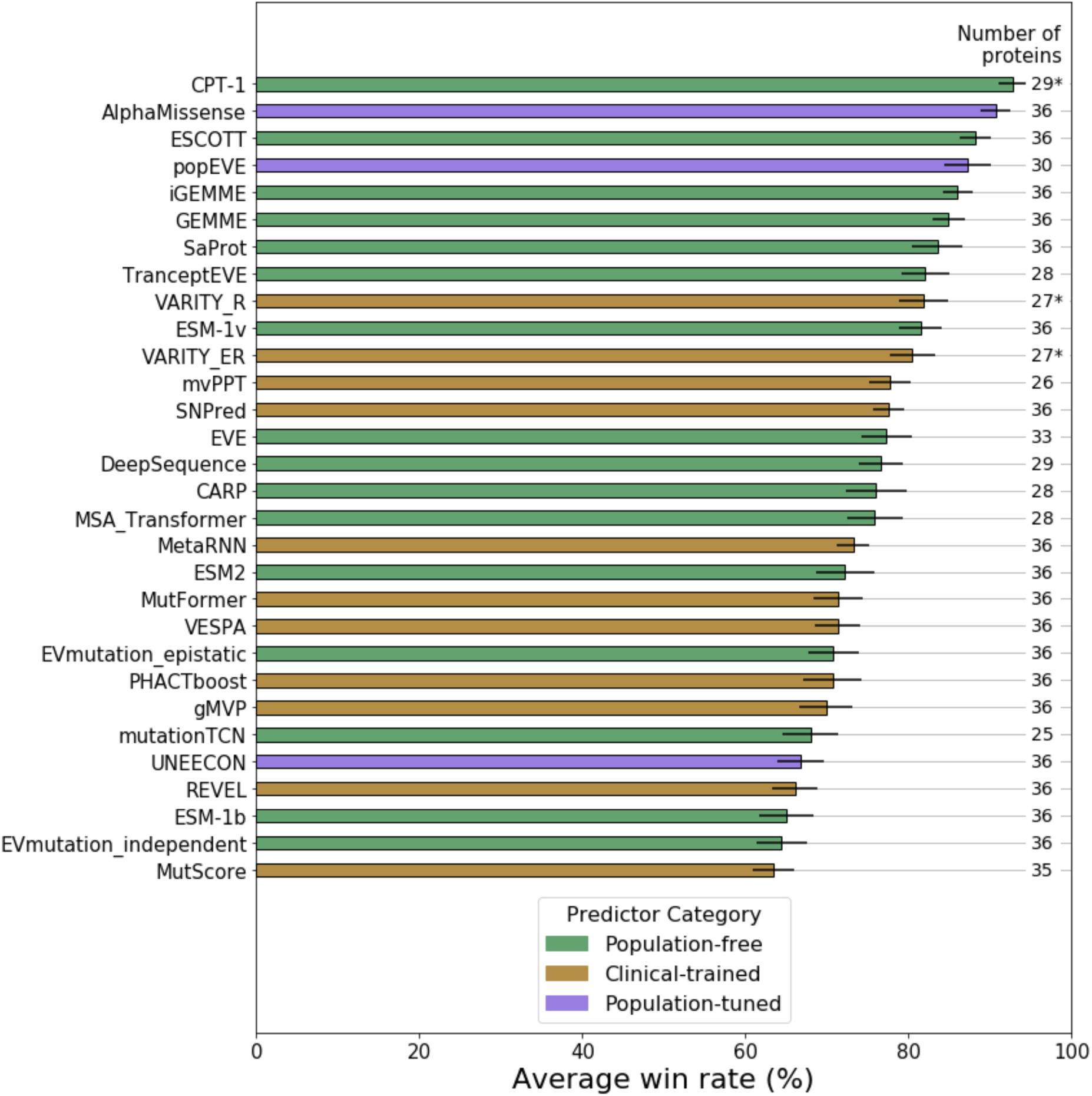
The top 30 out of 97 tested VEPs ranked based on performance against the DMS benchmark. VEPs are ranked according to their average win rate against all other VEPs in pairwise Spearman’s correlation comparisons across all DMS datasets. The number of proteins for which each VEP had scores included is indicated in the right column of the plot. Those indicated with * had some DMS datasets excluded to avoid circularity concerns. Error bars represent the standard deviation in the rank score across 1000 bootstrap permutations of the benchmarking DMS datasets. The full ranking of all VEPs and all pairwise win rates are available in Table S3, additional file 1.

The top-ranking VEP using this methodology is CPT-1 with a 92.8% overall average win rate and 66.2% average win rate against the other top 5 ranking methods. CPT-1 ranked significantly higher than every method in the bootstrap except AlphaMissense (p=0.169) (Table S4, additional file 1). CPT-1 combines both EVE [32] and ESM-1v [31] along with structural features from AlphaFold [36] and ProteinMPNN [37], and further conservation features. Importantly, although training of the model was carried out against five DMS datasets, they have all been excluded from the benchmarking of CPT-1, thus avoiding circularity concerns.

AlphaMissense performs only marginally worse overall than CPT-1 in this benchmark with a 90.7% overall average win rate and 65.3% average win rate against the other top 5 ranking methods. While coming second, it did not perform significantly better than either popEVE [38] (p=0.108) or ESCOTT [39] (p=0.094). AlphaMissense is a recently developed large language model with additional structural context based on the AlphaFold methodology, and fine-tuned on allele frequencies from humans and other primates. While the core of the model is unsupervised and population-free, the fine-tuning process using human variants necessitates its inclusion as a population-tuned predictor.

ESCOTT, iGEMME [39] and GEMME [40] are three closely related population-free predictors that rank 3^rd^, 5^th^ and 6^th^, respectively. GEMME is a relatively simple model, compared to the other top performers, based on epistasis between positions through evolution. GEMME also has lower computational requirements than comparably performing VEPs, and similar computational time to running a language model like ESM-1v. iGEMME is an optimised version of GEMME using deeper alignments and capable of efficiently handling larger proteins. ESCOTT is based on iGEMME and takes into account the likely structural context of mutated residues.

Two variants of EVE are also amongst the top predictors. popEVE ranks 4^th^ and is a hybrid of ESM-1v and EVE that also performs gene-level calibration on variants from UK Biobank, with the goal of making scores from different genes directly comparable. The usage of Biobank data by popEVE means that it is not necessarily subject to the same circularity concerns as clinical-trained VEPs, so we have classified it as population-tuned. TranceptEVE [41] ranks 8^th^ overall, and is a hybrid of Tranception [42] and EVE [32].

SaProt [43] is a unique protein language model, which leverages the foldseek tool [44] in order to encode structural context into the tokens provided to the model. This approach has allowed SaProt to out-perform all the ‘pure’ language models, ranking 7^th^. However, the hybrid language models – AlphaMissense, CPT-1 and popEVE – all rank higher.

VARITY_R ranked 9^th^ overall, and was the top-ranking clinical-trained predictor that was also included in our previous study. Interestingly, while VARITY_R previously ranked behind ESM-1v, EVE and DeepSequence, it has slightly outperformed them here. However, we also note that VARITY_R and VARITY_ER are compared to fewer DMS datasets than most other VEPs in this benchmark, necessitated by exclusion of DMS data on which it was directly trained.

The heatmap in Figure S3 highlights two small anomalies in the ranking. First, AlphaMissense out-performs CPT-1 on the DMS data, but CPT-1 ranks top overall as a result of it more consistently out-performing the other top predictors in pairwise analyses. Second, Wavenet [45] stands out in the heatmap with an unusual pattern due to its extreme heterogeneity in performance when tested against different proteins. For example, while it is the top performer in terms of Spearman’s correlation with the HRAS assay, it ranks 65^th^ overall due to its inconsistency.

It is likely that the methodology underlying the DMS assays this benchmark is based on can have a substantial impact on the results. To investigate this, we repeated the analysis, limiting the DMS studies to direct assays only, and repeated again with indirect assays. The CPT-1 model ranked top in both repeats, followed by AlphaMissense (Table S5, additional file 1). The VARITY predictors performed better when benchmarked on the indirect assays but the analysis is, in general, resistant to permutations and sub-setting of the DMS assays used to benchmark against.

Despite our efforts in producing a VEP-focussed benchmark, it is possible that the differential variant coverage of the different predictors could still lead to pairwise comparisons where the lower coverage VEP is favoured. A similar phenomenon was noted in the analysis of the CAGI6 Annotate-All-Missense challenge [46]. This might arise where the lower-coverage VEP provides no predictions for a region of a protein where it would otherwise perform poorly. To address this issue, we performed two additional analyses. First, the main reason for missing data among VEPs is the inability of some predictors to score multinucleotide variants, so we repeated the benchmark, restricting variants to only those possible through a single nucleotide change (Table S6, additional file 1). The rank scores only changed minimally, with the largest changes resulting in less than a 4-point difference in score (MSA Transformer [47]). Second, to address all missing predictions not due to SNV limitations, we repeated the analysis, filling-in all remaining missing predictions with the most benign score produced by each predictor. This is done on the assumption that most large prediction gaps should be due to poorly conserved protein regions and thus enriched in benign variation. The results of this analysis, shown in Table S7 (additional file 1), were a drop in the performance of SPRI by 6 points, and a relative increase in performance for VESPA [48], PANTHER, mvPPT [49] and TranceptEVE by about the same amount. Other than popEVE and TranceptEVE increasing in rank by 1 place each, the order of the top-ranking predictors is largely unaffected. These results help confirm the robustness of our benchmarking methodology.

### Performance of VEPs on clinical variant classification

The above DMS-based benchmarking of VEP performance might not be reflective of performance in pathogenicity prediction, which is the main practical application for which these methods are used. DMS assays are heterogeneous in their methodologies and in the meaning of their functional scores. This has been a common criticism of assessing VEP performance using functional assays [30,50] and it is very understandable. To what extent do our DMS-based rankings reflect utility for clinical variant classification?

Traditional assessment and comparison of VEPs has typically involved testing their discrimination between known pathogenic and known benign or putatively benign variants, often using datasets such as ClinVar [19] and gnomAD [20]. However, this can be extremely difficult to do in a fair manner for clinical-trained predictors. First, most supervised VEPs have been directly trained on pathogenic variants, so to compare performance, one needs to know the identities of all the variants used to train each predictor, and then find a set of variants not used by any of the tested VEPs. One also needs to ensure that no variants from the same positions as variants used in training, or even at homologous positions [51], are included in the test set.

An even stricter requirement for assessment of VEP performance for variants across different genes is that, in most instances, one should exclude any variants from the test set from any gene for which any variants were used in training, or from any homologues of these genes. That is, supervised VEPs should only be tested on different and non-homologous genes to any used in training, not just different variants. This is necessary to avoid gene-level circularity; otherwise, predictor performance will be inflated because models will learn to associate certain genes with pathogenicity, regardless of their ability to discriminate between variant effects within those genes [1,23]. Importantly, however, as long as performance assessment is carried out on a per-gene basis (*e.g.* the performance metric is calculated for each gene/protein separately), then it should generally be acceptable to test VEPs on genes on which they have been trained, as this avoids any risk of gene-level circularity. Alternately, variant labels can be balanced on a per-gene basis (*i.e.* the ratio of pathogenic to benign variants is equal in every gene) [46].

There are further concerns related to the identities of variants used as the negative class. The same requirement to not use variants used in training is equally true for these. However, a critical complication arises from the fact that many VEPs now incorporate human allele frequency information as a feature. This is severely problematic for the use of known benign variants (*e.g.* those classified as benign or likely benign in ClinVar) as the negative class. As allele frequency is routinely used in the classification of variants as benign [2], VEPs that includes allele frequency as a feature will likely suffer from circularity in these analyses. Even if allele frequency was not directly used in the clinical classification, common variants are simply more likely to be studied and receive a classification.

An alternative to using known benign mutations as the negative class is to use variants observed in the human population (*e.g.* taking all of those from gnomAD), which will mostly be very rare. We strongly advocate this approach for several reasons. First, it minimises the aforementioned circularity issue regarding the use of allele frequency to classify benign variants, although it does not completely eliminate it. Second, it is much more reflective of the actual clinical utility of variant effect predictions. The major challenge for clinicians is not in discriminating between common benign and rare pathogenic variants. Instead, it is in the much more difficult problem of distinguishing rare benign from rare pathogenic variants. Previous predictors, notably REVEL [52] and VARITY, have acknowledged this issue and tailored their models to the problem of rare variant identification. Finally, using rare population variants allows for much larger negative classes. In many disease genes, there are no or few variants classified as benign, severely limiting the number of genes for which reliable analyses can be performed.

Here, we have assessed the performance of VEPs in distinguishing between pathogenic and likely pathogenic ClinVar missense variants, and ‘putatively benign’ gnomAD v4 missense variants, taking all of those not classified as pathogenic. We recognise the limitation of this, in that there is likely to be a small proportion of as-of-yet unclassified pathogenic variants in our negative class, particularly in recessive genes and those with incomplete penetrance. Nevertheless, we believe that the advantages stated above far outweigh this issue. We generated predictions for 819 proteins that had at least 10 (likely) pathogenic and 10 putatively benign missense variants using 51 clinical-trained, 30 population-free and 6 population-tuned VEPs. It was necessary to exclude a few VEPs from this analysis because predictions were not available for enough proteins, particularly those where we obtained scores from ProteinGym. For each protein, we tested the discrimination between pathogenic and putatively benign for each VEP by calculating the area under the receiver operating characteristic curve (AUROC), which is a common measure of classifier performance that summarises the trade-off between true positive rate and false positive rate across different thresholds.

The full distribution of AUROC values for each predictor, sorted by median, is shown in Figure S4. However, for the same reasons discussed earlier in relation to the DMS ranking, this analysis has the potential to be confounded by the fact that not all VEPs provide scores for all possible variants.

Therefore, we applied the same pairwise ranking strategy as above, using AUROC as our comparison metric instead of Spearman’s correlation. Figure 3 shows the top 30 ranking predictors in terms of their performance in clinical variant classification according to this methodology. The full ranking of all available predictors is provided in Table S8, additional file 1.

**Figure 3.**
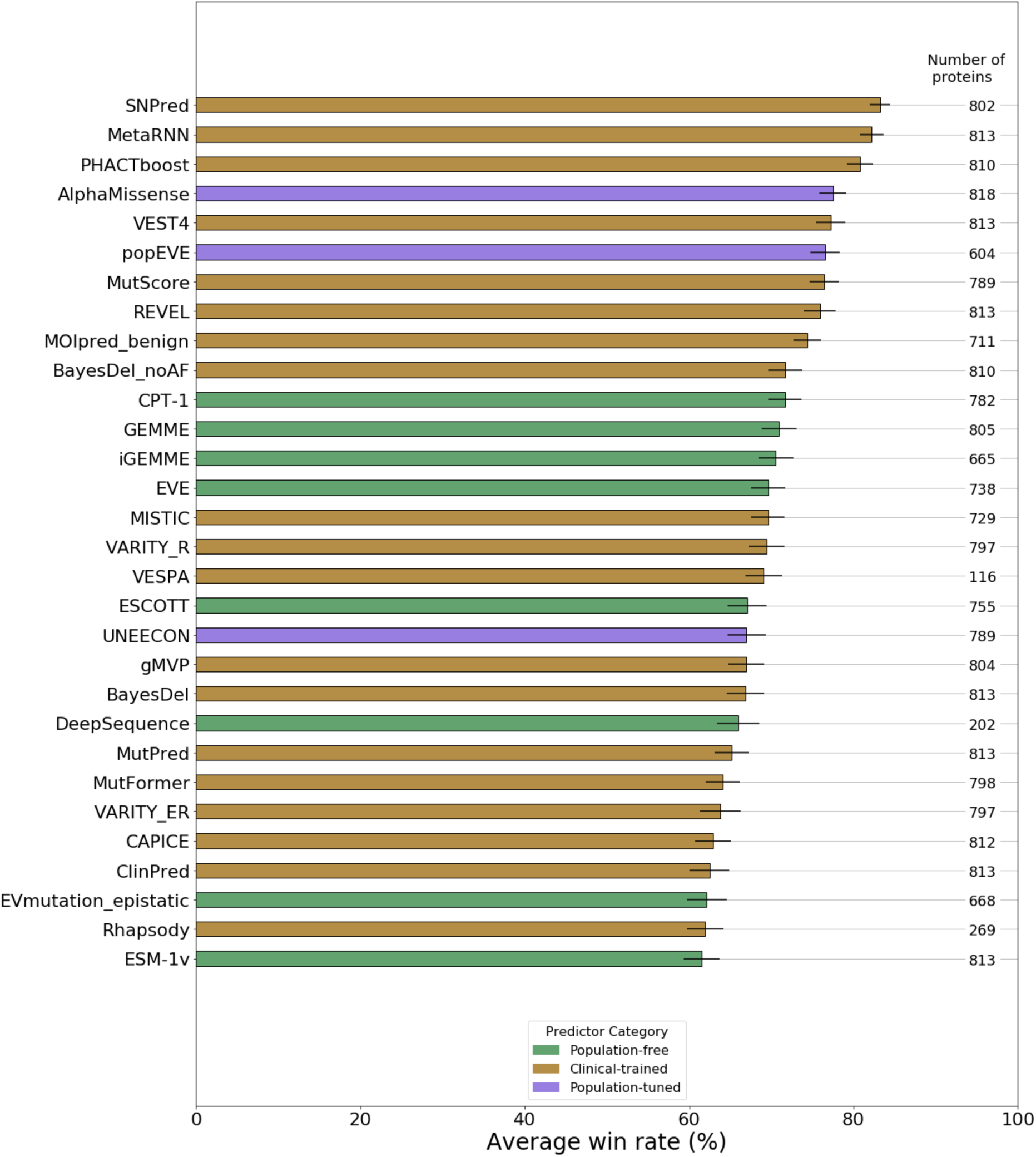
The top 30 out of 83 tested VEPs in terms of clinical variant classification performance. VEPs are ranked according to their average win rate against all other VEPs in pairwise AUROC comparisons across all human proteins with at least 10 pathogenic and 10 putatively benign missense variants. The number of proteins that met this condition for each predictor is indicated on the right of the plot. Some VEPs from the DMS benchmark could not be included here because predictions were not available for enough genes. Error bars represent the standard error across all comparisons with other VEPs. The full ranking of all VEPs and all pairwise win rates are available in Table S8, additional file 1.

At first glance, the rankings are strikingly different from the DMS-based analysis, with none of the top 10 ranking VEPs being population-free. SNPred [53] and MetaRNN [54] rank 1^st^ and 2^nd^, respectively, in contrast to the DMS benchmark, where they ranked 13^th^ and 18^th^ overall. It is likely that their performance here, as well as the performance of most clinical-trained VEPs, is highly inflated by circularity issues, as we made no effort to exclude variants used in training. Therefore, it is interesting to note that the top-ranking population-free VEP was CPT-1, the same as observed with DMS. The closely related GEMME, iGEMME and ESCOTT models show very similar performance, with ESCOTT ranking slightly higher in the DMS benchmark and GEMME/iGEMME ranking slightly higher in the clinical benchmark.

Area under the precision-recall curve (AUPRC) is an alternative performance metric to AUROC. Precision-recall is considered more reflective of many real-life classification scenarios where correct identification of a minority class is more important than that of a majority class. The disadvantage of precision-recall is that relative class sizes need to remain consistent across all models in order for the AUPRC scores to be comparable. Our use of pairwise comparisons essentially cancels out this disadvantage, allowing us to use AUPRC as an alternative to AUROC. Figure S5 ranks the predictors by pairwise analysis using AUPRC as the comparison metric. The overall rankings are very similar to the ROC-based ranking in Figure 3, but with 12 clinical-trained and population-tuned predictors exceeding the performance of the top population-free predictor.

To compare the two benchmarks, in Figure 4, we plot the win rate from the DMS analysis *vs* the win rate from the clinical variant analysis. If we consider only the population-free models, the Pearson correlation (r = 0.979) is striking. Relative performance on the DMS benchmark appears to be highly predictive of relative performance in clinical variant classification across the entire performance range. In contrast, for the clinical-trained models, the correlation is much lower (r = 0.841). While the Pearson correlation for the population-tuned VEPs is even higher (r=0.995), it is less significant due to the lack of predictors falling into this category (p=0.001). Moreover, the clinical-trained VEPs tend to show relatively increased performance on the clinical benchmark compared to the population-free VEPs. This almost certainly reflects varying levels of circularity contributing to performance in the clinical benchmark. It is likely that the extent to which clinical win rates are shifted to the right relative to the population-free VEPs can be considered as measure of how overfit the models are on our pathogenic and putatively benign variants. This is reinforced by the observation that the population-tuned VEPs are also right-shifted compared to the population-free methods but less-so than the clinical-trained VEPs, lending credence to the idea that they are intermediate to the two other groups in terms of circularity concerns.

**Figure 4.**
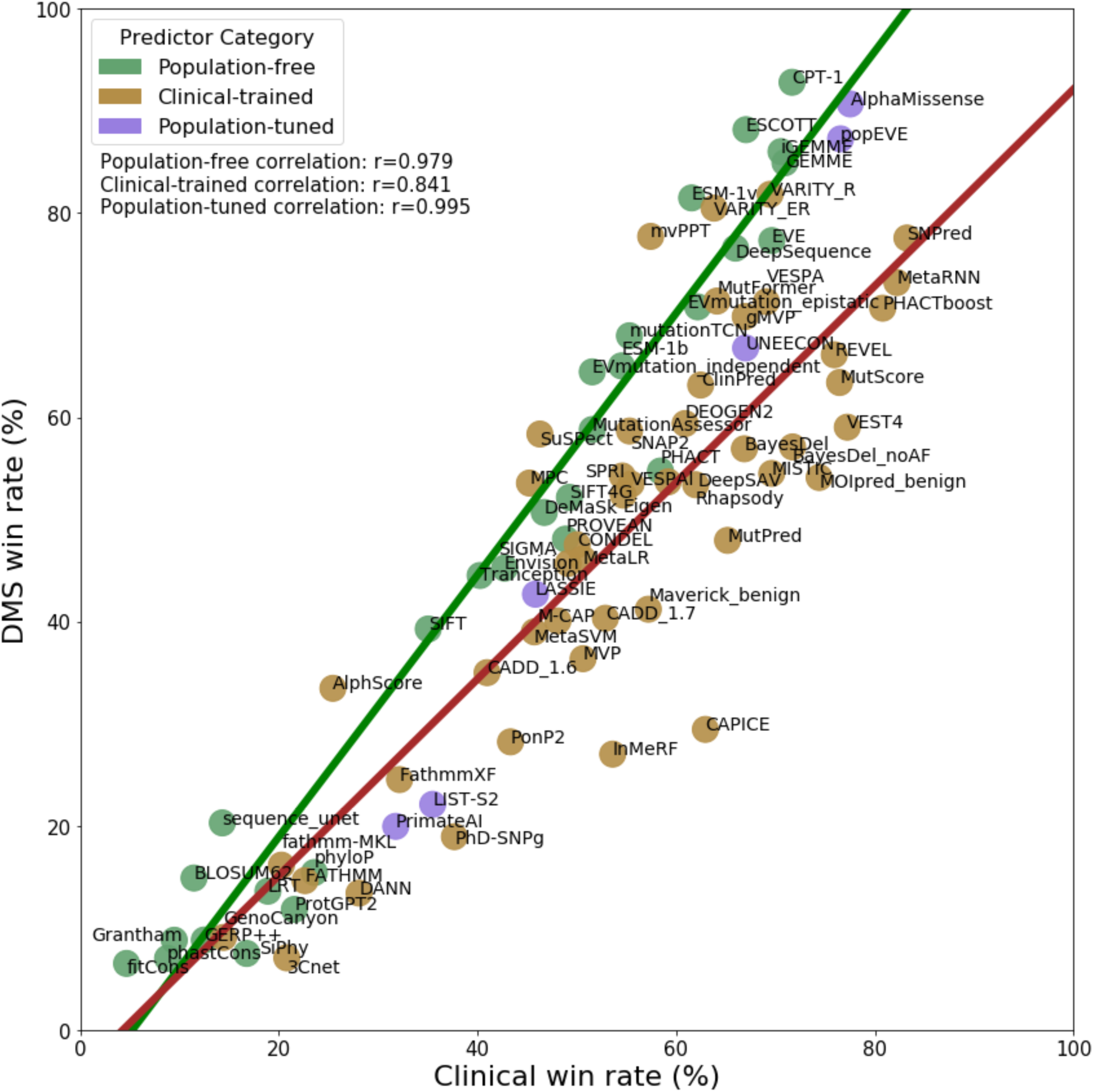
Strong correspondence in relative performance of VEPs on the DMS *vs* clinical benchmarks. Average pairwise win rates in the DMS *vs* clinical benchmarks are plotted. Population-free and population-tuned VEPs show extremely strong correlations (Pearson’s r = 0.979 and 0.995 respectively). In contrast, the clinical-trained VEPs show a much weaker correlation overall (r = 0.841). The tendency for some clinical-trained VEPs to show large rightward shifts, reflecting relatively increased performance on the clinical benchmark, is likely to be due to circularity due to training on variants and genes present in the pathogenic and putatively benign datasets.

While most of the clinical-trained VEPs show strong signs of circularity in their clinical variant classification performance, demonstrated by their right-shift in Figure 4 compared to the population-free methods, some appear to have much less or no bias. The two VARITY models fit perfectly with the population-free VEPs, possibly reflecting its innovative strategy to minimise training bias. mvPPT, SuSPect [55] and MPC [56], while ranking lower overall in both categories, also appear to show little bias.

The population-tuned predictors demonstrate some level of right-shift in Figure 4, relative to the population-free methods, although less than the majority of clinical-trained predictors. This indicates that there is likely some level of data circularity influencing their predictions on the clinical dataset, although it is not as severe as for the clinical-trained predictors. Both AlphaMissense and popEVE in particular are very close to the population-free trend. Our use of mostly rare variants as the putatively benign dataset should minimise any advantage to AlphaMissense from its population tuning. On the other hand, popEVE only uses population variants for scaling scores on a protein-level, to aid with cross-protein comparison of scores. In principle, this approach should not make the method vulnerable to variant-level circularity (although it could potentially be conflated by gene-level circularity in other cross-gene analyses). The remaining population-tuned methods are all closer to the clinical-trained trend.

### Practical considerations

An often-overlooked but extremely important aspect of VEPs is how easy they are for an end-user to obtain predictions. VEPs are typically made available through a combination of three different channels.

1. A web interface that allows access either to the VEP itself (*e.g.* SIFT [57], PolyPhen-2) or to a database of pre-calculated results (*e.g.* popEVE, VARITY).
2. A large compilation of pre-calculated results that usually cover either all canonical human protein positions in UniProt [58] or all human coding region non-synonymous single nucleotide variants in genome space.
3. The method itself is made available for installation by the end user.

Of these three channels, a web interface is the most convenient for looking up single variants of interest, although most such interfaces also offer the option to view all possible variants within a given protein as well. Downloadable databases of pre-calculated results are very useful for large-scale analyses (such as this one), but may be less useful for end users than a simple web interface for searching individual variants. If such a database is formatted in genome space, then specialised software such as Tabix [59] may be required to identify scores for variants of interest. Finally, installing and running the predictor offers the greatest degree of control over generation of the results such as the alignments and features used. However, many modern VEPs have high computational and time requirements or require significant technical knowledge to operate. We are unable to recommend such VEPs for typical day-to-day usage unless the data is also obtainable through a web interface or database.

As these are all important considerations for end users, in Table 3 we provide a summary of the top 15 VEPs from the Spearman’s correlation-based analysis in terms of how easy it is to obtain predictions, as well as links to any online interfaces, pre-calculated results or installable packages/repositories.

**Table 3.**
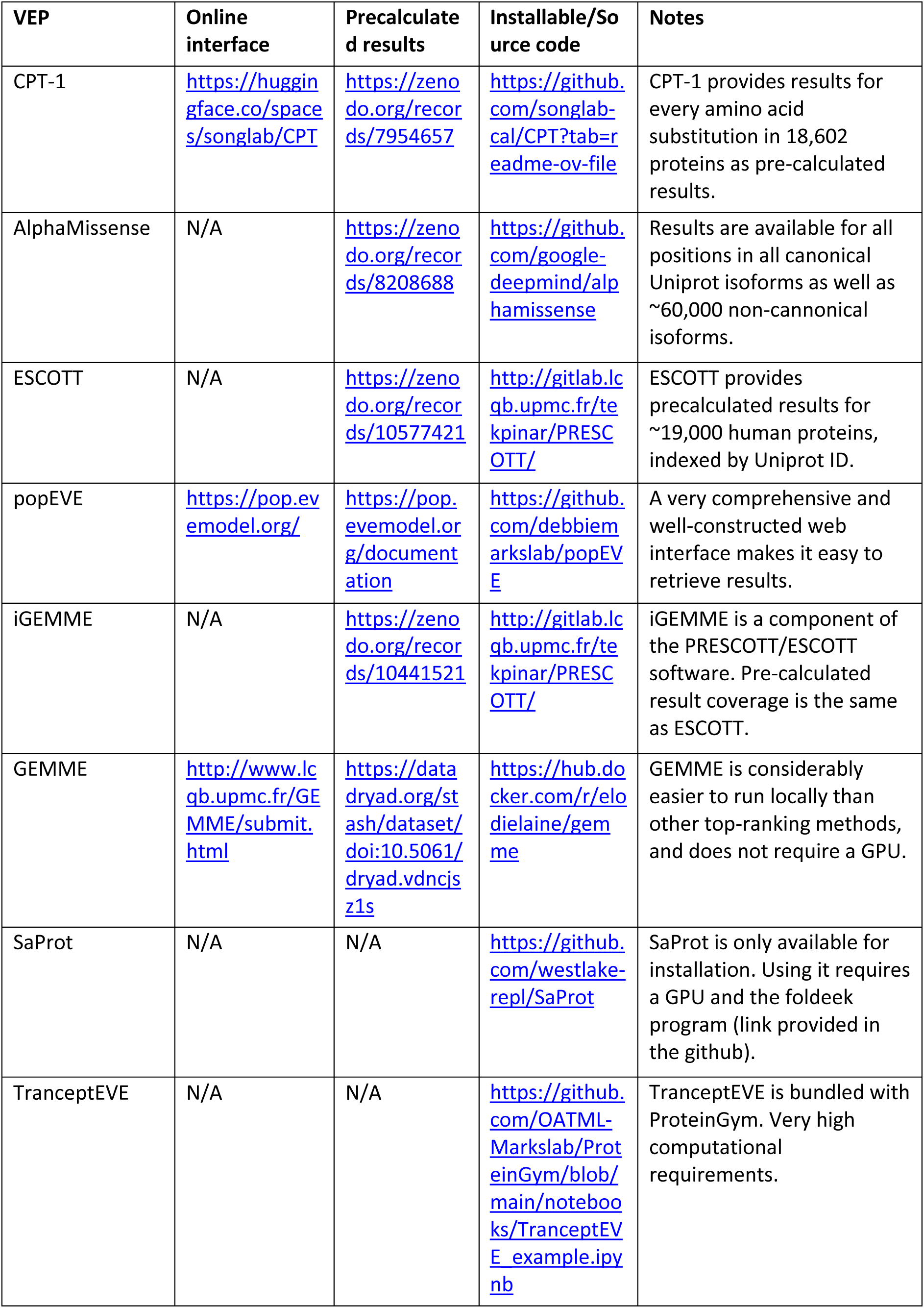
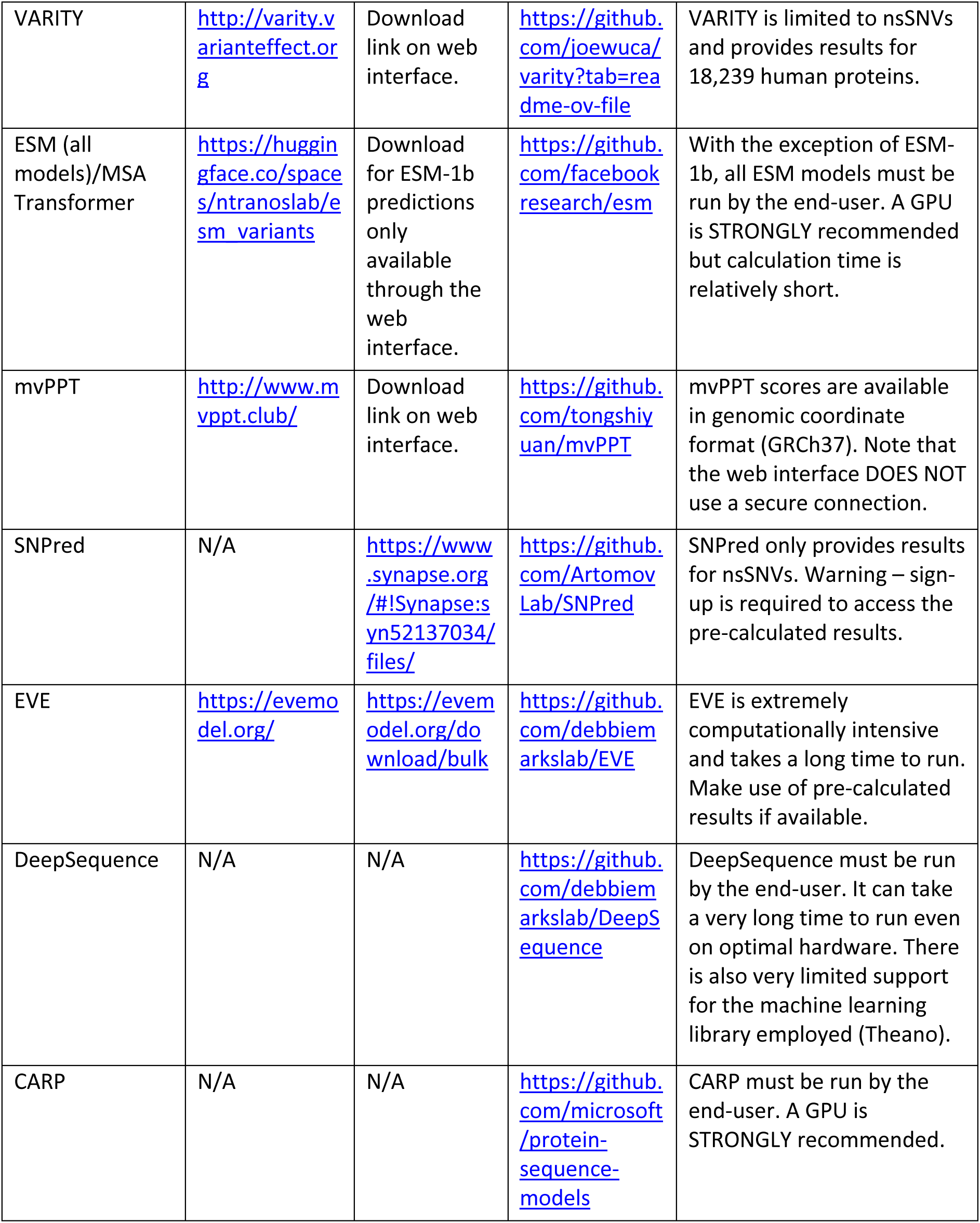
Summary of the top-ranking VEPs and the channels through which their results can be accessed.

## Discussion

Our benchmarking strategy very much relies on comparing performance across a large number of diverse DMS datasets compared to previous benchmarks. We have tried to avoid making judgements about the quality of individual DMS datasets or selecting them based on what we deem to be desirable phenotypes or experimental properties, other than excluding a small number based on irrelevance to disease mechanisms. Although it is possible (perhaps likely) that certain types of DMS experiments will be better for VEP benchmarking than others, we feel that our approach of taking as many datasets as possible minimises the potential for bias. Although different DMS datasets differ greatly in the extent to which they reflect clinical phenotypes, they generally should show at least some relationship to fitness and thus, in general, algorithms that are better at predicting variants effects on fitness or pathogenicity should tend to show a stronger correlation with experimental measurements. The fact that we see such a strong correspondence between the relative ranking of VEPs across these diverse DMS datasets, and in the clinical classification of variants, strongly supports the utility of this approach. Importantly, however, our strategy requires comparing performance across numerous datasets, as we observe large variability in the ‘winner’ from dataset to dataset. Thus, any attempts to judge performance with datasets from one or a small number of functional assays are unlikely to yield very informative results.

This analysis improves upon our previous benchmark in three important ways. First, the additions of new VEPs and DMS datasets allowed considerable expansion of the benchmark and allows us to assess how the state-of-the-art methods perform when compared to established ones in common use. Second, our switch from a DMS-focussed ranking method to a pairwise, VEP-focussed ranking method allows a much fairer comparison between predictors that fail to make predictions in certain protein regions. The robustness of this ranking method is demonstrated in Table S7 (additional file 1) where the exclusion of non-SNV variants and filling-in of prediction gaps resulted in very minor changes in predictor ranking. Finally, our findings that VEP rankings on DMS datasets is strongly correlated with their performance on clinical datasets and the differences between the three classes of VEPs gives additional validity to our methodology and demonstrates the impact of data circularity.

From the differences between the VEP categories in Figure 4, It is clear that much of the tendency for clinical-trained VEPs to perform relatively better on the clinical benchmark is due to data circularity. However, one could still argue that there is some aspect of the relatively better performance that comes from these VEPs learning some aspect of clinical pathogenicity that is not present in the population-free models. We think this is unlikely. For example, the strategy used by the clinical-based VARITY went to great lengths to minimise circularity issues in its training process, and it is highly consistent with the population-free VEPs in its relative performance on DMS *vs* clinical benchmarks. AlphaMissense was not trained for pathogenicity, but even with its inclusion of allele frequencies, it is also fairly similar to the population-free methods. Finally, CPT-1, without any training on human pathogenic or population variants, outperforms 38/48 tested clinical-trained VEPs on the clinical benchmark, demonstrating how effective population-free methods can be on their own.

One related concern that is very difficult to address is optimisation against DMS datasets present in our benchmark. While we have excluded datasets directly used in training for the evaluation of certain predictors, it is possible that methods may have been optimised against DMS data without direct training. For example, ESM-1v was not trained on DMS data, but it was selected out of multiple possible models based on its correlation with DMS data [31]. Possibly this is reflected in the fact that it is slightly “left-shifted” in Figure 4, showing modestly better performance on the DMS benchmark relative to the clinical benchmark. As DMS and other functional assays are increasingly used to assess performance, VEP developers will inevitably target these benchmarks and optimise for performance against them. However, currently there is little indication that DMS inclusion in VEP training or optimisation has had an impact on this benchmark, and the few methods trained directly with DMS data have proven to be relatively resistant to bias. We hope that this methodology can continue to be used to benchmark future predictors in a bias-free manner.

Other recent studies have also attempted to assess the performance of state-of-the-art VEPs using alternate strategies. Tabet *et al* tested the ability of 24 different VEPs to infer human traits from the UK Biobank and All of Us cohorts [60]. While this task is distinct from predicting pathogenicity, it should be largely free of any circularity concerns. Interestingly, their overall rankings are broadly similar to what we observe here. While many of our top-ranking VEPs were not included in their study, the three top methods they identified based on UK Biobank data, AlphaMissense, ESM-1v, and VARITY, all ranked higher in our DMS benchmark than any other methods they included, and overall, there is a strong correlation (r = 0.93) between our DMS win rate and their number of traits identified (Fig. S6).

The CAGI Annotate-All-Missense challenge presents an extremely valuable study, testing the performance of 60 VEPs in discriminating between 10456 pathogenic and benign missense variants [46]. They avoid the issue of variant-level (type 1) circularity by only used variants deposited in ClinVar or HGMD after a specific cutoff date, and only considering predictions submitted before this date, or from methods that are not trained on clinical variants. They also explore the fact that methods that directly use allele frequency essentially have an unfair advantage when classifying benign variants, showing that the performance of these VEPs is much worse when considering rarer variants. While their main analysis does not account for gene-level circularity, they also present a gene-label balanced analysis, where they only consider equal numbers of pathogenic and benign variants from each gene. While this greatly reduces the size of their dataset, to 2140 variants from 504 genes, it should entirely overcome the issue of gene-level circularity. If we exclude those VEPs that directly use allele frequency, we see some similar results in their gene-balanced analysis compared to our DMS-based ranking. Specifically, their top three methods are AlphaMissesne, ESM-1v and EVE, which also perform better in our DMS benchmark than any of the other VEPs included in their study.

An interesting observation from our latest DMS benchmark is that the three top-ranking methods, CPT-1, AlphaMissense and ESCOTT, all use some level of protein structural information. Previously we had noted that there was no tendency for structure-aware models to perform better than those that use sequences only [15], so this represents a potentially notable advance in VEP development. One possibility is that, in the past, the large majority of performance gains have been obtained through improved elucidation of the evolutionary signal, and so the much smaller impact of structure was negligible. This is compounded by the fact that most structure-based VEPs assume that pathogenic variants will be structurally damaging and ignore non-loss-of-function effects [61], and that, previously, structure models were only available for a minority of human proteins. Thus, given the recent availability of computational structural models for all human proteins [36,62], the inclusion of structural information may now becoming more important for varaiant effect prediction.

Given the remarkable performance of population-free VEPs, we think that not directly including human clinical or population variants in models is the safest strategy for variant effect prediction. Given the desire to increase the role of computational predictions in making clinical diagnoses, it is important to minimise the potential for “double counting”, *e.g.* according to ACMG/AMP guidelines [2]. If allele frequency, or knowledge of other classified variants at the same position, have been utilised by the model, then the computational prediction cannot be considered as independent evidence. In contrast, population-free VEPs should be truly independent from the other pathogenic or benign classification criteria.

Although we believe that our relative rankings of VEP performance are reliable, a major remaining problem is still in the interpretation of their outputs. For example, how should a clinician interpret a high VEP score for making a genetic diagnosis? Recent work has attempted to establish thresholds for using variant effect scores as stronger diagnostic evidence [63]. This is a potentially powerful approach, but it does have limitation. Given the radically different performance of VEPs across different genes, it is not clear that the same thresholds for evidence levels will be appropriate for different genes [64–66]. Furthermore, this work focused primarily on clinical-trained methods, and calibrating these VEPs using known pathogenic and benign variants is likely to overstate the confidence with which pathogenic or benign evidence can be assigned due to the same circularity-related issues discussed here, especially for methods that directly utilise allele frequency.

Overall, it is clear the variant effect prediction field is moving very fast. Along with other members of the Atlas of Variant Effects Alliance, we recently released a set of guidelines and recommendations for developers of novel VEPs, many of which related to improving the sharing and independent assessment of methods [23]. In addition, we strongly encourage researchers to deposit new DMS datasets in MaveDB [18]. Making methods, predictions and DMS data freely and easily available will improve future DMS-based benchmarking. Finally, we note that, while missense variant effect prediction is reaching a level of maturity, far more work remains to be done on non-missense coding variants and on non-coding variants, both in terms of methods development and benchmarking. We hope the lessons we have learned here will prove valuable for this.

## Conclusions

In this study we have used functional data from 36 diverse DMS experiments to benchmark the relative performance of 97 VEPs while greatly reducing the potential for bias compared to traditional benchmarks. Our pairwise comparison methodology is robust to both the datasets employed and missing predictions. We demonstrate the data circularity issue with benchmarks based on clinical data, and provide recommendations for general-use VEPs. We expect the scale of this type of benchmark to expand in scope over time, although training and optimisation of VEPs against DMS data may hinder such efforts in the future.

## Methods

### ClinVar and gnomAD data

We obtained ClinVar data from ClinVar on 06/08/2024. We then filtered the data by retaining only missense variants labelled as ‘Pathogenic’, ‘Likely pathogenic’ and ‘Pathogenic/Likely pathogenic’. We then removed all entries with a 0* review status (no assertion criteria) and all entries with conflicting interpretations of pathogenicity.

Our gnomAD dataset was obtained from gnomAD version 4.1. We retained all missense variants that passed either the gnomes or exomes FILTER criteria. We refer to this dataset as ‘putatively benign’ and while it certainly contains some recessive or low-penetrance variants at low frequency, represents the distribution of variants in a healthy population.

### DMS datasets

Starting with the 26 DMS datasets from out previous benchmark, we excluded all variants present within our ClinVar and gnomAD datasets, then retained only datasets that scored at least 1000 amino acid variants. We also excluded datasets that measured antibody binding. This resulted in the exclusion of *BRCA1* (insufficient variants remaining), *CCR5* and *CXCR4* (antibody binding). We identified a further 13 datasets that also met our inclusion criteria. These new datasets were primarily obtained from MaveDB [18], but also by searching published works. One dataset (*GCH1*) came from an unpublished study with permission of the authors.

### VEPs

The dbNSFP database, version 4.2 [67] served as a source for 27 VEPs. The remaining 70 VEPs were either run locally on the University of Edinburgh high performance computing cluster (EDDIE), downloaded as pre-calculated results, obtained via a web interface or obtained for a limited subset of mutations/proteins from the ProteinGym website. Table S2 (additional file 1) provides a full list of the source used to obtain predictions from each VEP.

### Spearman’s correlation and rank score

Spearman’s correlation was calculated between datasets using the stats.spearmanr() function of the scipy python package version 1.5.4 on Python version 3.6.8.

To calculate the correlation-based rank score displayed in Figure 2 and Table S3 (additional file 1), for each protein the absolute Spearman’s correlations between the selected DMS dataset and every pair of VEPs was calculated using only variants where the DMS and two VEPs all have available scores. The VEP that obtained the highest correlation in each pairwise comparison earned one point, while the VEP with the lower correlation earned none. The win rate of every VEP over every other VEP was then calculated across all proteins by dividing the number of wins by the number of times that particular pair of VEPs were tested. The final rank score was calculated by averaging the win rate of each VEP against all other VEPs.

### AUROC and AUPRC

The area under the receiver operator characteristic curve (AUROC) was calculated using the metrics.roc_auc_score() function of the sklearn python package, while the area under the precision-recall curve (AUPRC) was calculated using the metrics.average_precision_score() function of the sklearn python package version 0.18.1. To maintain consistency between class labels, predictors that assigned low scores as pathogenic and high as benign needed to be inverted. This was done by deducting the scores from 1.

The rankings in Figure 3 were calculated by comparing the AUROC between every pair of predictors using only variants shared between them. The predictor with the highest AUC was awarded one point. The win rate of every VEP against every other VEP was then calculated across all proteins by dividing the number of wins by the total number of times that particular pair of VEPs were tested. The final rank score was calculated by averaging the win rate of each VEP against all other VEPs. The same procedure was used to generate the AUPRC-based ranking in Figure S4, but with precision-recall instead of ROC.

## Authors contributions

The initial idea for the study was conceived by JM. BL performed all data analysis, produced the figures and took the lead in writing the manuscript. The final version of the manuscript was produced with significant input from JM. Both authors read and approved the final manuscript.

## Acknowledgements

This work has made use of the resources provided by the Edinburgh Compute and Data Facility (ECDF) (http://www.ecdf.ed.ac.uk/). The authors would also like to thank Zebinisa Mirakbarova for identifying several VEPs benchmarked in this study, Mihaly Badonyi for collating and mapping the data from ClinVar and gnomAD and Lukas Gerasimavicius for proofreading.

## Supplemental Figures

**Figure S1.**
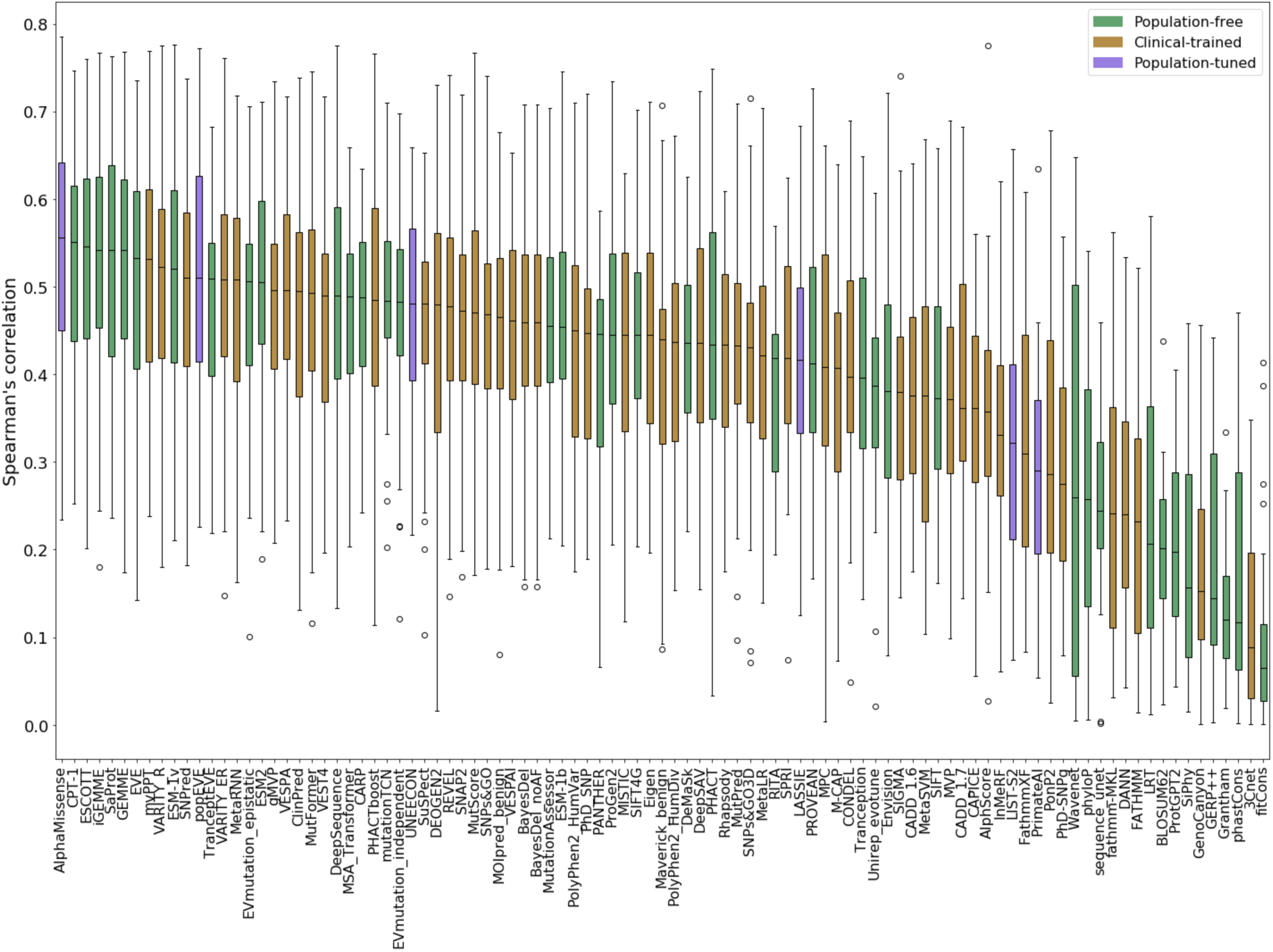
Distribution of Spearman’s correlations of different VEPs across different DMS datasets. This is effectively the same data presented in Figure 1, but grouped by VEP instead of by protein. Correlations of different VEPs across different DMS datasets. Note that not all VEPs output scores for all DMS datasets, or all variants from individual datasets, so we can not necessarily directly compare correlations; thus motivating the pairwise ranking approach in Figure 2.

**Figure S2.**
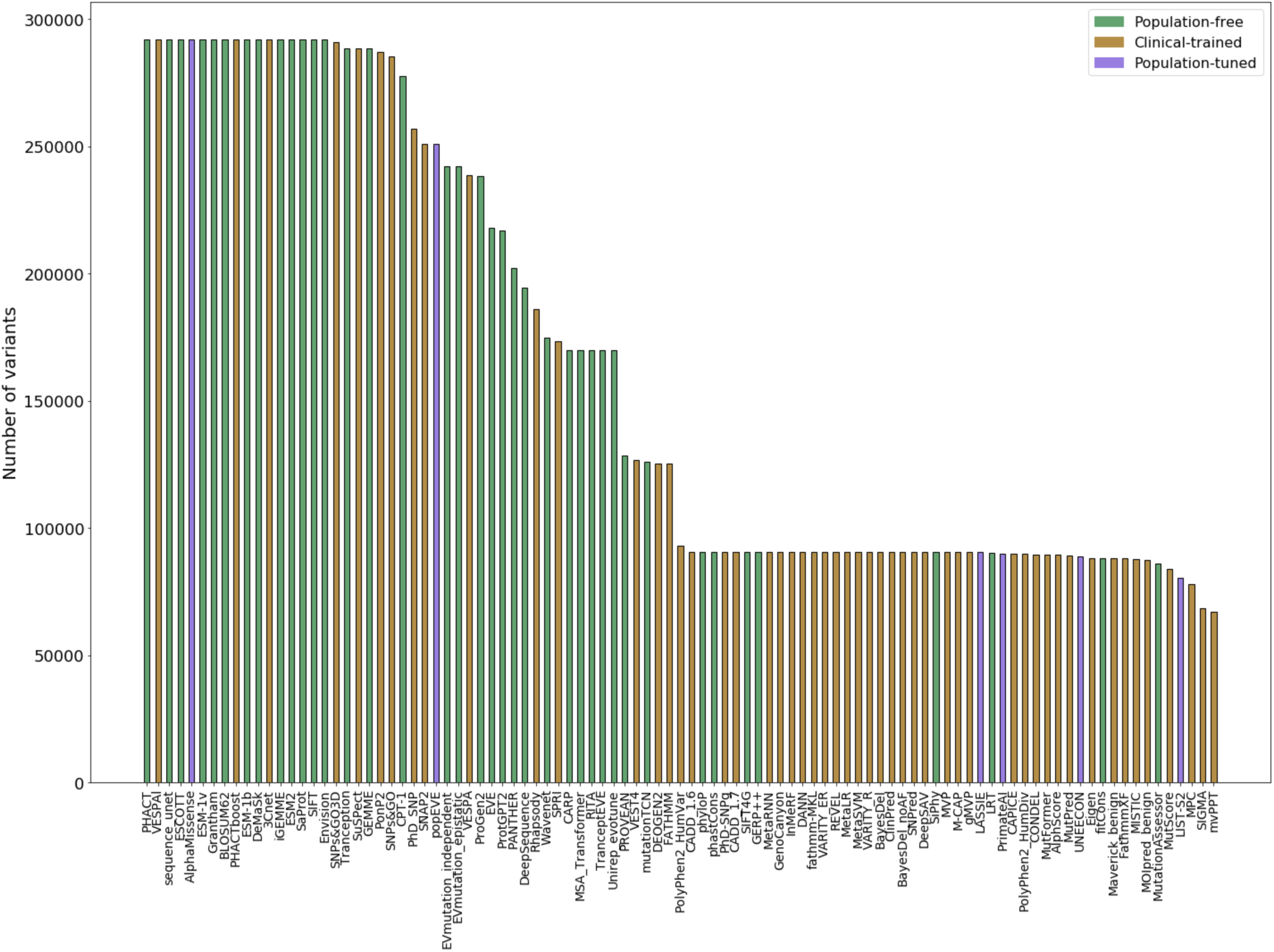
Variant coverage of amino acid substitutions by VEPs. The number of variants scored within our dataset by each VEP is shown in this figure. This does not necessarily truly demonstrate the utility of some VEPs such as PolyPhen-2, which are capable of scoring more variants, but are not available for mass-download.

**Figure S3.**
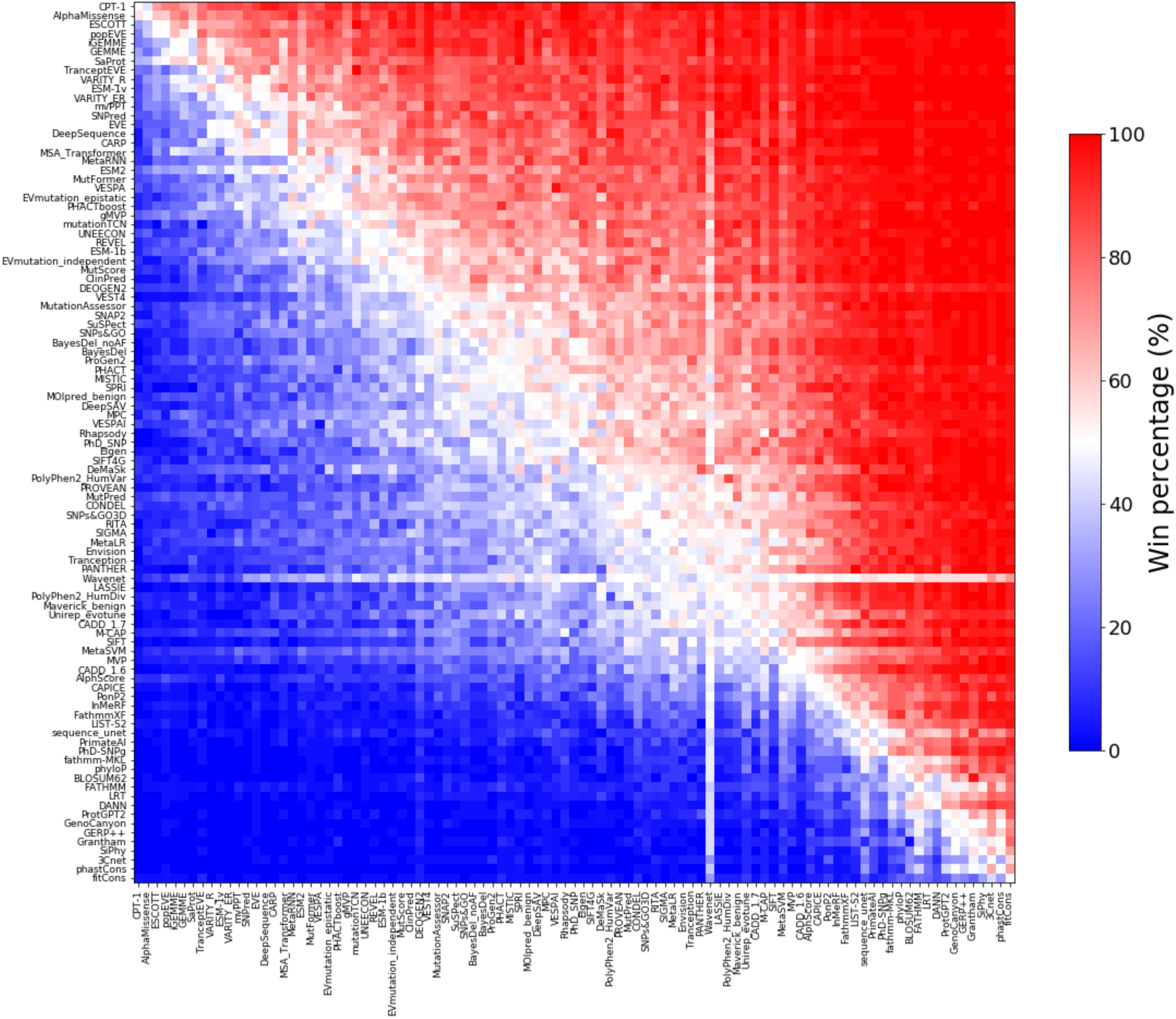
Heatmap of VEP vs VEP win rates across DMS datasets. For each pair of VEPs, we compare their Spearman’s correlations with each DMS dataset, considering only variants for which each VEP has a prediction available. The win rate represents the percentage of proteins where the first VEP shows a higher correlation than the second. For example, red values indicate the VEP on the y-axis outperforms the VEP on the x-axis in terms of correlation with DMS measurements across most proteins. The average win rate in Figure 2 represents the mean win rate of each VEP against all other VEPs.

**Figure S4.**
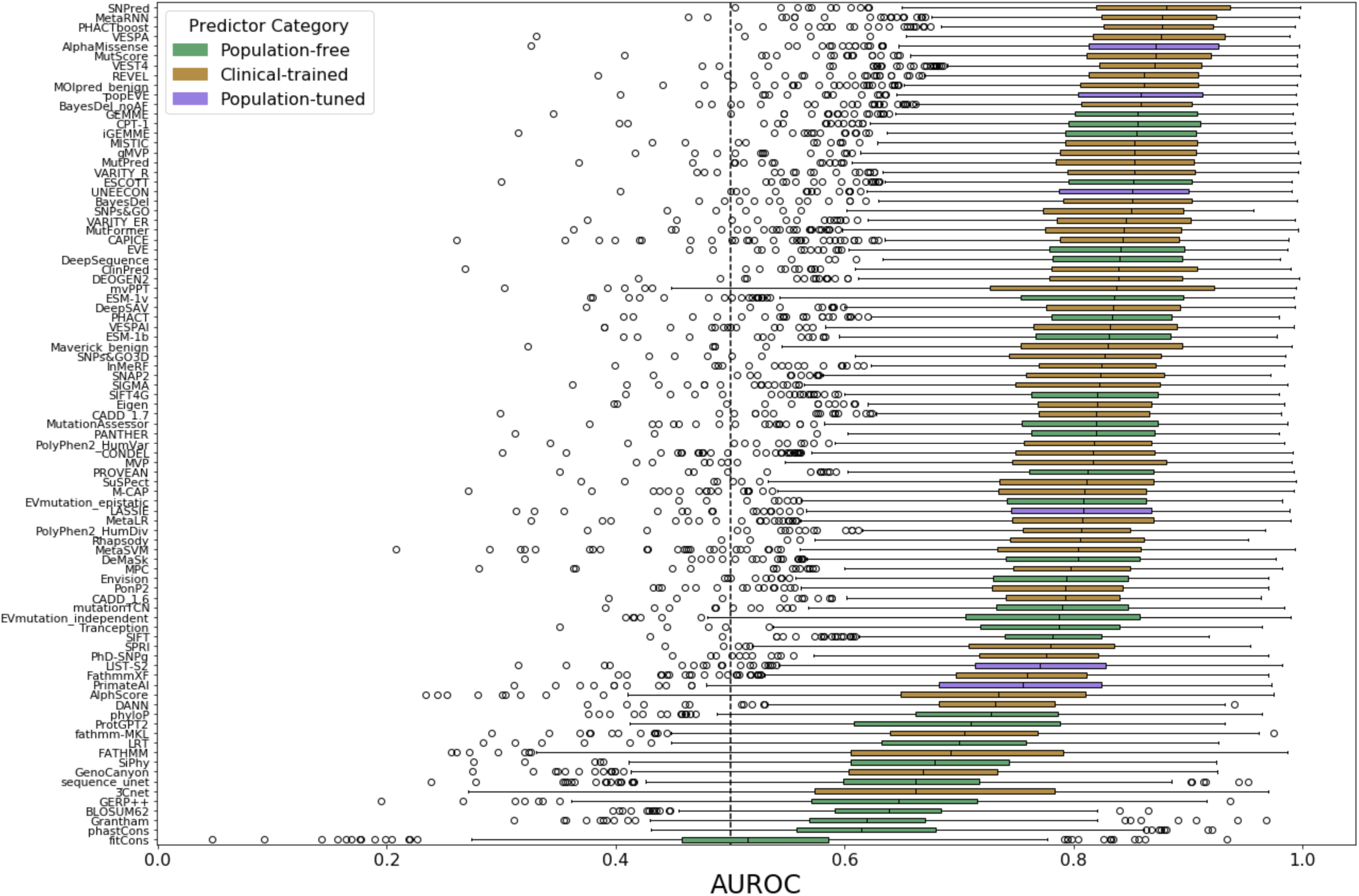
Distribution of AUROC representing discrimination between pathogenic and putatively benign missense variants for each VEP across different human protein-coding genes. As for the DMS comparison, not all VEPs output predictions for all proteins or all variants. Thus, the pairwise ranking represents a better reflection of relative performance.

**Figure S5.**
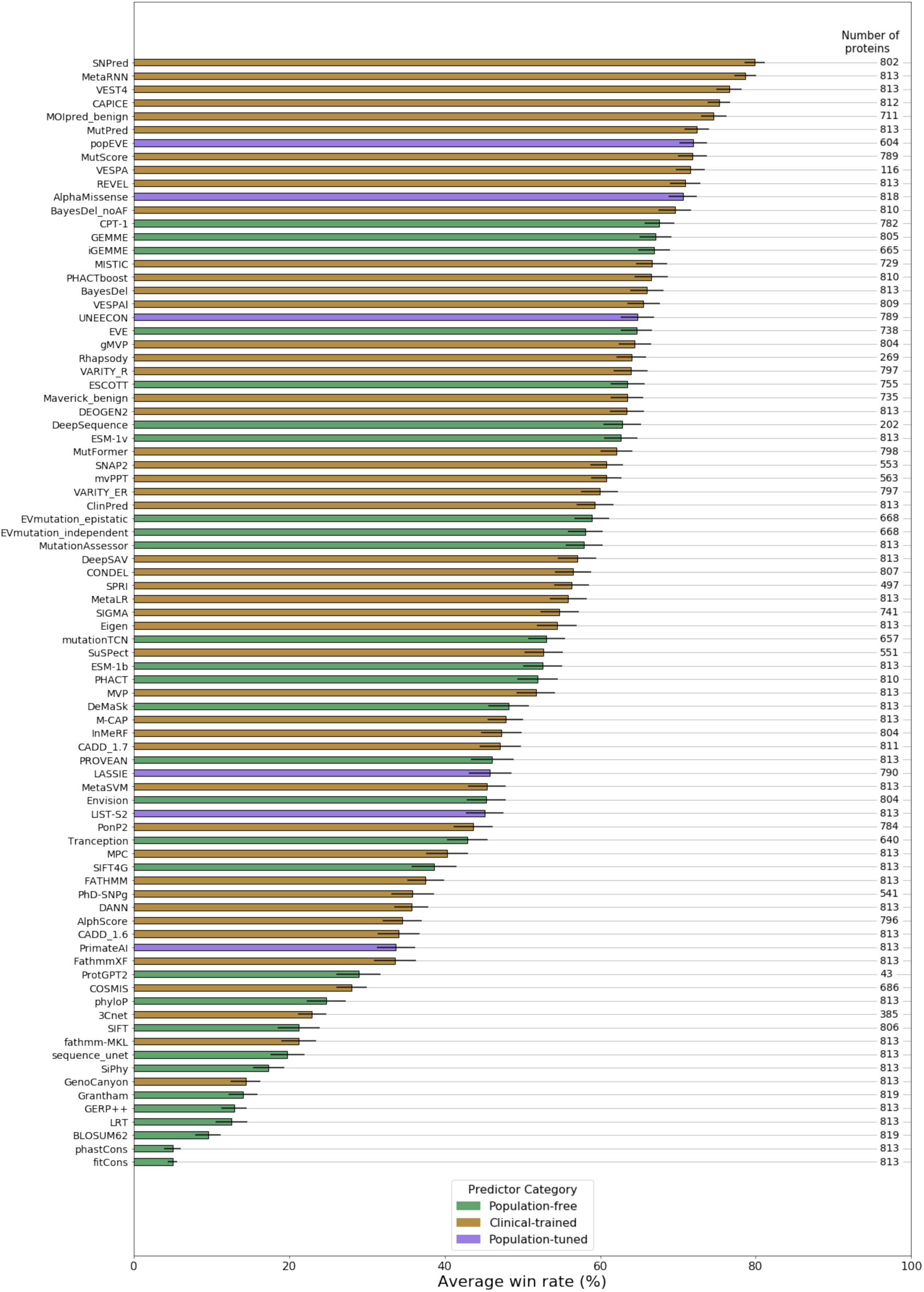
Ranking of VEPs on clinical variant classification using AUPRC. This utilises the same strategy as in Figure 3, but with precision-recall instead of receiver operating characteristic curves used in pairwise comparisons. Error bars represent the standard error across all comparisons with other VEPs.

## Additional Files

**Additional file 1.** Excel spreadsheet (.xlsx format). Contains Supplemental Tables 1-8 and full references for all VEPs and DMS datasets in the study.

